# Dopamine Abundance Uncouples Neurodegeneration and Lifespan in a *C. elegans* Model of Parkinson’s Disease

**DOI:** 10.64898/2026.07.04.736516

**Authors:** Corey W. Willicott, Tyler J. Altman, Logan C. Kimble, Laura A. Berkowitz, Guy A. Caldwell, Kim A. Caldwell

## Abstract

The neuropathology of Parkinson’s disease is characterized by α-synuclein (α-syn) aggregation and dopaminergic (DAergic) neurodegeneration. While neuronal loss in *C. elegans* α-syn-induced neurodegeneration models is temporally age-dependent, prior research indicates it is uncoupled from the organismal aging process. Here we examined transgenic *C. elegans* expressing human A53T α-syn in DAergic neurons to determine the impact of localized DA metabolism on both neurodegeneration and organismal lifespan. Increasing endogenous DA levels through overexpression of tyrosine hydroxylase (CAT-2) exacerbated A53T-induced DAergic degeneration, whereas DA depletion via Δ*cat-2* mutation rescued neuronal survival. By mutating a DA-interaction motif within α-syn, neurodegeneration was rendered insensitive to DA manipulation, thus confirming a structural basis for in vivo toxicity. We identified a DA-α-syn interaction that acts as a common upstream bridge whereby localized stress induces physiological responses in *C. elegans*. Genetically, this biochemical interaction acts as a pleiotropic trigger driving two compartmentalized responses: localized DAergic neurodegeneration via oxidative stress, and organism-wide, TFEB/*hlh-30*-dependent proteostatic remodeling that extends lifespan. Modulating autophagy, without exacerbating DA-mediated oxidative stress, represents a promising strategy to preserve adaptive systemic remodeling while limiting targeted neuronal damage.

## Introduction

Parkinson’s disease (PD) is defined neuropathologically by the progressive loss of dopaminergic (DAergic) neurons and the accumulation of intracellular inclusions enriched in α-synuclein (α-syn) (Spillantini et al., 1997; Braak et al., 2003). A critical, unresolved paradox in PD pathology is the relationship between the onset of neurodegeneration and organismal aging. While neuronal loss in *C. elegans* Parkinson’s models increases with time, it is an *age-dependent*, rather than an *aging-dependent*, phenomenon (Apfeld and Fontana, 2017); the loss of individual neurons is entirely uncoupled from organismal lifespan. The biochemical mechanism that permits this targeted neurodegeneration to proceed independently of the systemic aging process remains unknown.

DA is intrinsically reactive and readily undergoes oxidation, generating quinones and reactive oxygen species capable of modifying proteins and perturbing cellular homeostasis (Hastings, 2009; Burbulla et al., 2017). α-Syn is a direct biochemical target of DA; DA-modified α-syn species exhibit altered aggregation kinetics and stabilization of soluble oligomeric intermediates (Conway et al., 2001; Norris et al., 2005; Mazzulli et al., 2007). Familial PD-associated mutations in α-syn, such as A53T, accelerate oligomerization and enhance proteotoxic stress (Polymeropoulos et al., 1997; Lázaro et al., 2014). Notably, DA-α-syn interactions depend on residues within the C-terminal region of α-syn, suggesting that structural coupling between DA and α-syn conformation can directly modulate toxicity (Norris et al., 2005; Mor et al., 2017).

Beyond its role in neuronal vulnerability, DA signaling influences organismal physiology and aging. In *Caenorhabditis elegans* (*C. elegans*), DA regulates locomotor behavior, metabolic state, and stress responsiveness (Sawin et al., 2000; Chase and Koelle, 2007). Longevity in *C. elegans* is tightly linked to proteostasis and autophagy pathways, including regulation by the TFEB ortholog HLH-30, which coordinates lysosomal biogenesis and autophagic gene expression and is required for multiple lifespan-extending paradigms (Lapierre et al., 2013; Madeo et al., 2015).

Previous work demonstrated that DAergic expression of A53T α-syn in *C. elegans* induces progressive neuronal dysfunction and degeneration, and that mutation of a putative DA-interaction domain attenuates this toxicity (Mor et al., 2017; Mor et al., 2019). We hypothesize that this DA-α-syn biochemical interaction is the molecular driver that uncouples localized neurodegenerative vulnerability from systemic adaptive stress remodeling. To address this, DA biosynthesis was genetically increased or reduced in animals expressing either A53T α-syn or a DA-interaction-deficient A53T α-syn variant. Neurodegeneration, functional behavioral deficits, stress resistance, lifespan, and autophagy dependence were systematically examined to determine whether DA-α-syn coupling acts as a molecular mediator balancing targeted proteotoxic vulnerability with adaptive organismal longevity.

## Results

### DA levels modulate neurodegeneration

To visualize dopaminergic (DAergic) neurons in vivo, we utilized the integrated transgenic *C. elegans* strain BY250 [*vtIs7* (P*_dat-1_::*GFP)] (Nass et al., 2002), which maintains its full complement of six anterior DAergic neurons [2 ADE (anterior deirid); 4 CEPS (cephalic sensory)] (Figure 1C) as hermaphrodite animals age (Figure 1A, D). To determine how endogenous DA levels impact neuronal survival, we modulated endogenous DA production in *C. elegans*. Overexpression of the *C. elegans* ortholog of tyrosine hydroxylase, CAT-2, in the DAergic neurons {UA57 [*baIs4* (P*_dat-1_*::*cat-2*; P*_dat-1_*::GFP)]} caused DAergic neurodegeneration [(Cao et al., 2005); (Figure 1A, E)].

**Figure 1.**
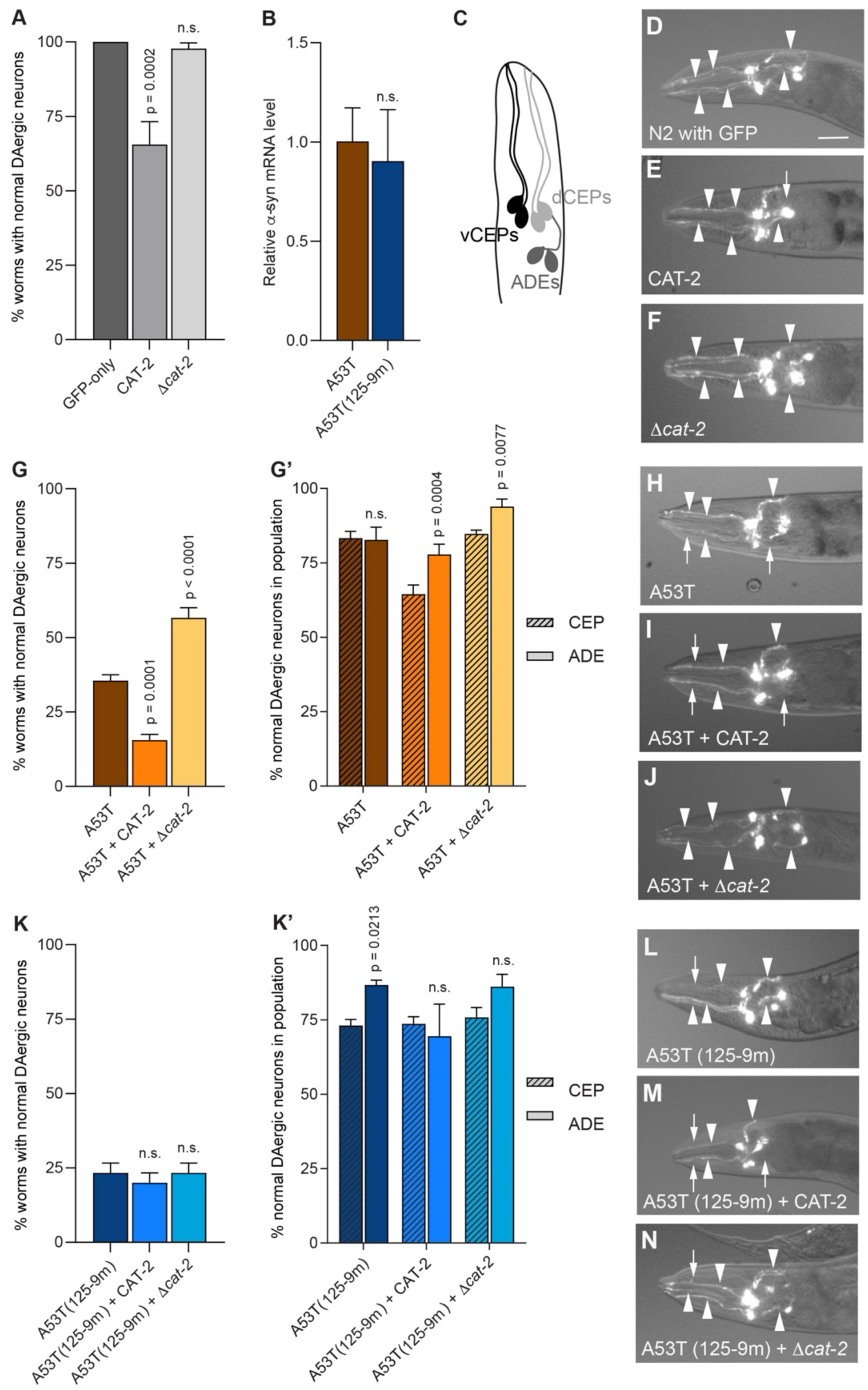
**α-Synuclein A53T DAergic neurodegeneration is modulated by DA perturbation**. **A**. Percentage of worms with normal DAergic neurons on day 7 post-hatching in GFP-only, CAT-2 overexpression, and Δ*cat-2* backgrounds. **B**. Relative α-syn mRNA levels measured by RT-qPCR in A53T and A53T(125-9m) strains. Values represent pooled RNA (n=120 worms per genotype per replicate) with three biological replicates and three technical replicates per genotype. A Student’s *t*-test was performed for statistical analysis. Expression level of α-syn was normalized to A53T animals. **C**. Schematic representation of the six anterior DAergic neurons (4 CEP, 2 ADE). **D - F**. Representative fluorescent micrographs of day 7 animals lacking α-syn. **G**. Percentage of normal DAergic neurons in A53T, A53T + CAT-2, and A53T + Δ*cat-2* animals, with **G’** breaking down survival by CEP versus ADE neuron class in day 7 animals. **H - J**. Representative micrographs of the A53T variant strains. **K**. Percentage of normal DAergic neurons in A53T(125-9m) variant strains, with **K’** breaking down survival by CEP versus ADE neuron class in day 7 animals. **L - N**. Representative micrographs of the A53T(125-9m) variant strains. Values represent mean ± S.D. n=30 worms per genotype per replicate, three independent replicates. One-way ANOVA with Dunnett’s post hoc test was used for comparisons. Representative fluorescent micrographs identify intact DAergic neurons with arrowheads and missing DAergic neurons with arrowheads. Scale bar, 20μm.

Conversely, the *cat-2(n4547)* allele has a 1010 bp deletion and is considered a near-null mutant because it leads to very low DA levels (Omura et al., 2012). It was crossed into BY250 (P*_dat-1_*::GFP) to create the strain UA502 {P*_dat-1_*::GFP [*vtIs7*] V; *cat-2(n4574)*}, which displayed no effect on DAergic neuron survival (Figure 1A, F).

To explore potential impacts of DA levels on α-syn-induced neurodegeneration, we created transgenic worm strains expressing the A53T α-syn mutation. Previously, mutation of the C-terminus in α-syn was reported to abolish the interaction between α-syn and DA, as well as inhibit DA-mediated stabilization of oligomers in vitro (Norris et al., 2005; Mazzulli et al., 2007; Herrera et al., 2008). This interaction is reversible and dependent on residue ; and the Y_125_EMPS_129_ motif in the C-terminus of α-syn (Conway et al., 2001; Norris et al., 2005; Mazzulli et al., 2006). We previously reported the generation of a transgenic *C. elegans* line where A53T α-syn was engineered to contain a substitution where amino acids Y_125_EMPS_129_ were changed to F_125_AAFA_129_ [α-syn A53T(125-9m)] (Mor et al., 2017). *C. elegans* transmitting stable transgenes of either α-syn A53T or α-syn A53T(125-9m) constructs were examined for evidence of changes in neurodegeneration. We observed DAergic neurodegeneration was more robust in α-syn A53T compared to α-syn A53T(125-9m) animals (Mor et al., 2017). Together with parallel mammalian studies, this provided the first in vivo evidence of a direct neurotoxic correlation between α-syn structure and DA levels (Mor et al., 2017; Mor et al., 2019).

These previously reported transgenic A53T-variant strains harbored stably expressing transgenes, which could display fluctuating phenotypes due to mosaicism (Frokjaer-Jensen et al., 2008). Therefore, to more definitively examine the role of DA on neurodegeneration in the A53T background, we generated *C. elegans* strains containing chromosomally integrated transgenes expressing either human α-syn A53T or α-syn A53T(125-9m). These were UA372 {P*_dat-1_*::α-syn(A53T), P*_unc-54_*::tdTomato[*baIs54*]} and UA373 {[P*_dat-1_*::α-syn A53T(125-129m)], P*_unc-54_*::tdTomato[*baIs55*]}, respectively. Here, we refer to these new chromosomally integrated lines as either A53T or A53T(125-9m) for clarity. A comparison of α-syn expression by RT-qPCR revealed no significant difference in mRNA levels between A53T and A53T(125-9m) (Figure 1B).

To examine DAergic neurodegeneration, A53T and A53T(125-9m) were crossed to BY250 (P*_dat-1_*::GFP) to create strains UA526 and UA530, respectively. Expression of A53T resulted in significant DAergic degeneration of neuronal cell bodies and processes where only 36% of the population at day 7 displayed normal DAergic neurons (Figure 1G, H). A53T(125-9m) demonstrated similar neurodegenerative trends whereby only 23% of the population at day 7 exhibited normal DAergic neurons (Figure 1K, L). These results contrasted with GFP-only expressing animals, which did not display neurodegeneration at day 7 and were 100% normal (Figure 1A, D). A more detailed analysis of DAergic neurodegeneration in A53T(125-9m) animals revealed that the pair of ADE neurons were more protected from degeneration than the two bilaterally symmetrical pairs of CEP neurons (p = 0.0213) (Figure 1K’). There was no significant difference between CEP and ADE neuroprotection in the A53T worms (Figure 1G’).

To explore how α-syn-induced neurodegeneration is impacted by altered DA levels, we crossed A53T and A53T(125-9m) animals with CAT-2 tyrosine hydroxylase-overexpressing animals to create strains UA525 and UA529, respectively. A significant increase in DAergic neurodegeneration was observed within the population of A53T + CAT-2 animals compared to A53T alone (16% vs. 36%; p=.0001) (Figure 1G). The ADE neurons were more protected from degeneration than the CEP neurons in the A53T + CAT-2 animals (p = 0.0004) (Figure 1G’, I). In contrast, overexpressing CAT-2 in A53T(125-9m) *C. elegans* did not alter DAergic neurodegeneration within animal populations (Figure 1K, M). Individual DAergic neuron types (ADE and CEP) were also not significantly impacted by CAT-2 overexpression (Figure 1K’). These results agreed with our previously reported findings using stable A53T and A53T(125-9m) transgenic lines (Mor et al., 2017), verifying that our newly integrated strains recapitulated the neurodegenerative patterns.

Next, we crossed the integrated A53T and A53T(125-9m) strains with Δ*cat-2* null mutant animals. As shown in Figure 1A, the *cat-2(n4547)* allele had no effect on DAergic neuron survival in GFP-only animals. However, in the presence of A53T, the loss of DA resulted in DAergic neuroprotection within populations of A53T + Δ*cat-2* animals (57% vs. 36%; p<.0001). (Figure 1G). In these animals, the ADE neurons were more protected from degeneration than the CEP neurons (p = 0.0077) (Figure 1G’, J). The *cat-2(n4547)* allele did not alter DAergic neurodegeneration within A53T(125-9m) animal populations (Figure 1K, N). Similarly, DAergic neurodegeneration in the ADE and CEP were not significantly impacted by the *cat-2* null allele (Figure 1K’).

### Functional deficits precede neurodegeneration in A53T overexpressing animals

Because functional impairments can precede overt neurodegeneration in α-syn models, DAergic circuit performance was assessed prior to structural decline. To connect our observations in Figure 1 we conducted a functional behavioral assay termed the basal slowing response (BSR). The BSR assay is performed on day 3 worms, to gain insight into neuronal dysfunction before α-syn-induced DAergic neurodegeneration is apparent and connect DAergic neuron survival to physiological outcomes (Martinez et al., 2017; Gaeta et al., 2022).

In *C. elegans,* DAergic neurons are mechanosensory; they mediate a motor circuit that is used to sense and respond to changes in the environment by modulating locomotive behavior. Healthy wildtype (N2) animals slow their locomotion rate when they encounter a bacterial lawn; this basal slowing response requires DA (Sawin et al., 2000). When DA is absent, animals are unable to sense this physical stimulus and fail to slow down. By calculating the ratio of speed off food to speed on food, we can determine how the BSR is altered compared to N2, which has a standardized BSR efficiency of 100% (Figure 2A). An efficiency of less than 100% indicates a smaller difference between the off/on food speeds and, therefore, defective BSR. Analysis of Δ*cat-2* mutant animals compared to N2 revealed a deficit in BSR efficiency (34%; p=0.0006). Application of 2 mM exogenous DA restored BSR to ∼100% (p=0.0005) (Figure 2A).

**Figure 2.**
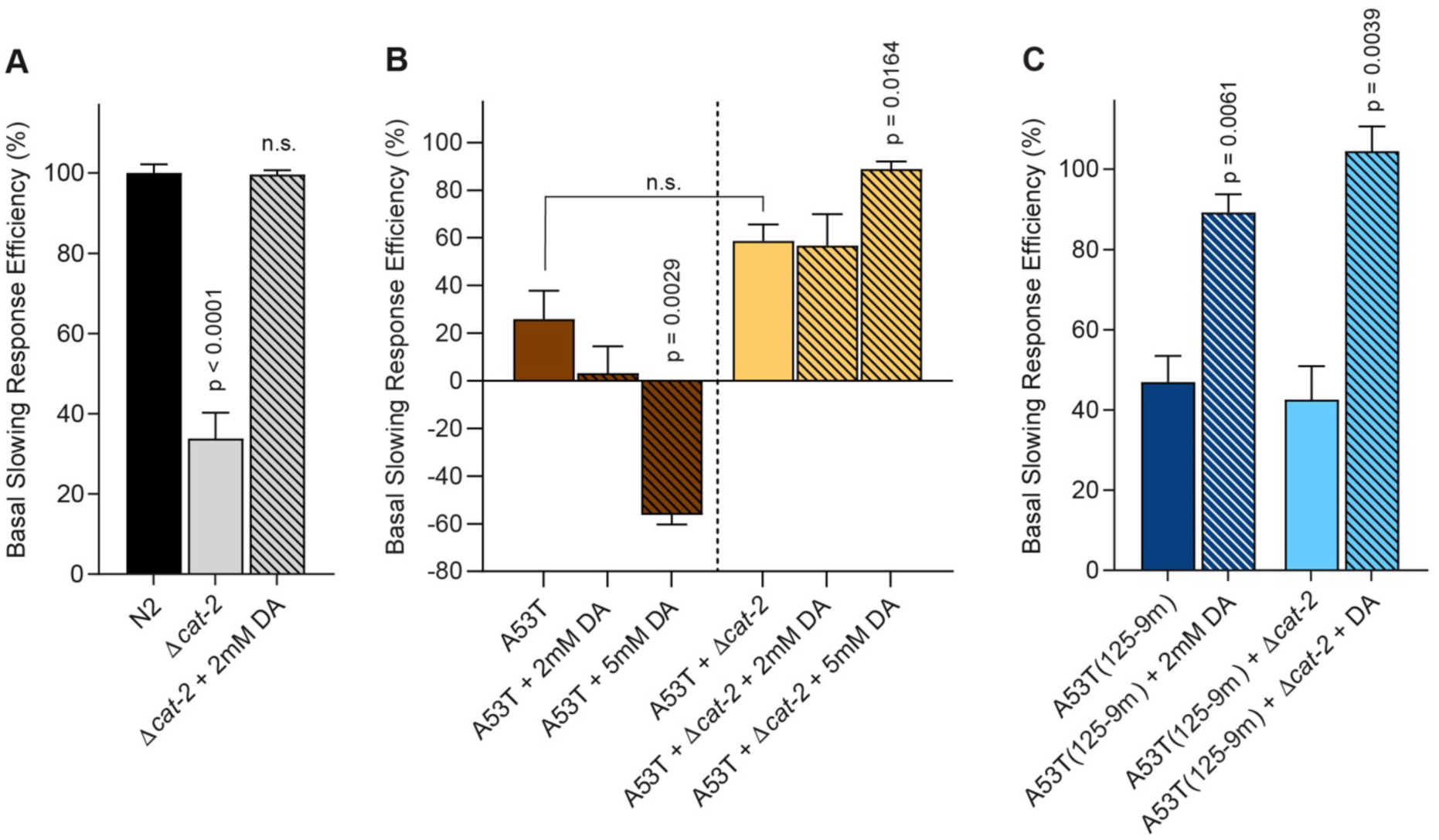
**Defects in basal slowing response highlight vulnerability of A53T DAergic neurons when exposed to exogenous DA**. Basal slowing response (BSR) efficiency assay normalized to N2. **A**. Δ*cat-2* mutants synthesize significantly less DA than N2 worms and serve as positive control. BSR can be rescued in these animals by treatment with 2mM exogenous DA. **B**. A53T overexpressing animals in an otherwise normal background (brown bars) or in the Δ*cat-2* mutant background (yellow bars) exhibit a defective BSR. Treating A53T animals (brown striped bars) with 2mM exogenous DA does not impact BSR while 5mM significantly decreased BSR. In the A53T + Δ*cat-2* animals, 2mM DA is non-significant but 5 mM significantly protected animals. **C**. A53T(125-9M) ± Δ*cat-2* animals displayed BSR defects that can be rescued with exogenous DA treatment (blue bars). Worms were assayed on day 3 post-hatching. Exogenous DA was applied for one hour prior to analysis. Values represent mean ± S.E.M. n=10 worms per genotype per replicate, with three independent replicates. One-way ANOVA with Dunnett’s post hoc analysis was used to compare within each group.

Although the BSR allows for earlier detection of degenerative dysfunction in DA neurotransmission, unpublished data from our lab found that the hypersensitivity of the BSR can result in it being altered when GFP is expressed in the DAergic neurons. This artifact was eliminated here by examining A53T and A53T(125-9m) strains that did not express GFP (UA372 and UA373). Compared to the 100% BSR efficiency of N2, A53T displayed a significant shortage, with a 26% BSR (p = 0.0037) (Figure 2A vs. B).

Exogenous DA treatment was used in lieu of the CAT-2 overexpressing strain because it has GFP as a co-injection marker. When A53T was treated with 2 mM DA no significant impact on the BSR was observed (Figure 2B). To further explore the negative trend, A53T worms were exposed to 5 mM exogenous DA. This concentration resulted significantly decreased BSR compared to A53T without exogenous DA (p = 0.0029) (Figure 2B).

The neuroprotective A53T + Δ*cat-2* animals had an improved, albeit non-significant, BSR compared to A53T (p = 0.0761) (Figure 2B). These animals were still deficient in BSR efficiency compared to wildtype (N2) animals, with a BSR efficiency of 59% (Figures 2B). When they were exposed to 2 mM DA, there was no significant change in BSR efficiency. However, increasing DA to 5 mM uncovered a rescuing effect for A53T + Δ*cat-2* animals, whereby BSR efficiency improved from 59% to 89% (p = 0.0164) (Figure 2B).

Next, A53T(125-9m) and A53T(125-9m) + Δ*cat-2* were examined in the BSR assay. Both genetic backgrounds had similar defective BSRs with values of ∼45%. Exogenous DA treatment resulted in a significant restoration of BSR for A53T(125-9m) to 89% efficiency (p=0.0061) and A53T(125-9m) + Δ*cat-2* to 104% efficiency (p=0.0039) (Figure 2C).

### Oxidative stress resistance in A53T animals

To determine whether DA-α-syn coupling alters systemic stress responsiveness, mitochondrial oxidative challenges were examined across age. Much like humans, *C. elegans* display an age-associated decline in both generalized movement and recovery from oxidative challenges (Bansal et al., 2015; Hahm et al., 2015; Banse et al., 2024). Therefore, to uncover any deficiencies in energetics associated with the A53T-DA interaction, all nematode strains were exposed to sodium azide (Figure 3A-C) or paraquat (Figure 3D-F), as these chemicals are known to induce mitochondrial ROS formation.

**Figure 3.**
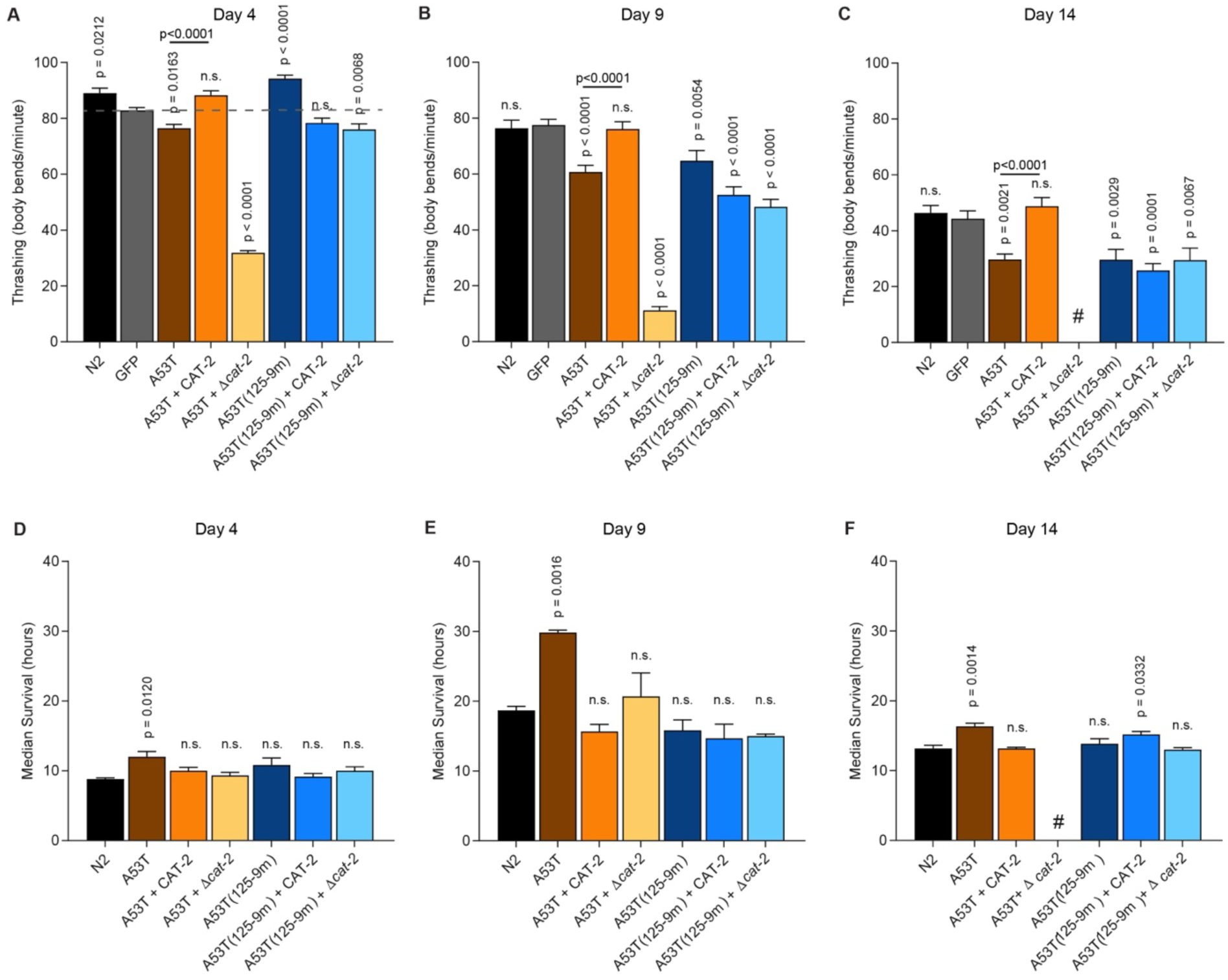
**The A53T-DA interaction drives a differential response to oxidative stressors**. **A - C**. Thrashing rates (body bends/minute) of animals on days 4, 9, and 14 post-hatching following an 8-hour exposure to 100 µM sodium azide. A53T animals display an energetic vulnerability that is rescued by CAT-2 overexpression. # indicates insufficient healthy A53T + Δ*cat-2* animals available for day 14 analysis. **D - F**. Median survival times of animals on days 4, 9, and 14 post-hatching following exposure to 40 mM paraquat on day of analysis. A53T animals exhibit a systemic survival advantage compared to N2 controls, which is abolished by altering DA levels (CAT-2 overexpression or Δ*cat-2*). Values represent mean + S.E.M. n=10 animals per genotype per replicate, three independent replicates. One-way ANOVA with Dunnett’s post hoc analysis was used to compare strains to their respective controls (GFP for azide; N2 for paraquat).

Every five days a population of each genotype were exposed to 100µM sodium azide-containing plates for four hours. Following the exposure, a well-characterized neurobehavioral analysis (thrashing in liquid) was performed. Across all strains, aging reduced the rate of thrashing, as demonstrated by the lower number of body bends/minute on days nine and 14 compared to day four (Figure 3A-C). On the first day of analysis (day 4), A53T(125-9m) animals displayed significantly better thrashing than GFP only animals (p <0.0001) while A53T animals were significantly worse (p = 0.0163) (Figure 3A). The dichotomy in thrashing behavior disappeared on days nine and 14 (Figure 3B, C), suggesting that the fortified stress resistance associated with A53T(125-9m) was not sustainable over time, as animals aged.

A53T *C. elegans* required sufficient DA levels to overcome sodium azide associated stress whereby both A53T and A53T + Δ*cat-2* animals demonstrated significantly worse thrashing rates from the control, GFP-only, animals (Figures 3A-C). In contrast, the A53T + CAT-2 animals were rescued at all timepoints (p<0.0001 for days four, nine, and 14) (Figure 3A-C). In this exposure paradigm, the A53T-DA interaction is associated with an energetic deficiency that is rescued by overexpression of CAT-2.

While sodium azide exposure revealed that the A53T-DA interaction drives an acute energetic vulnerability in motor function, we next asked whether this same interaction might paradoxically prime *C. elegans* to survive a systemic oxidative challenge. To investigate this, we examined the strains for overall survival following exposure to paraquat. In this assay, every five a population of animals was placed in a 40 mM paraquat solution and checked for life every hour until death. In general, we found that middle-aged animals (day nine) had better resistance to paraquat-induced stress when compared to counterparts on days four and 14 (Figure 3D-F). On days four, nine, and 14, we identified statistically significant increased survival times for A53T-overexpressing animals compared to N2 animals (Figure 3D-F). The A53T + Δ*cat-2* animal population did not survive until d14 to complete the last day of this assay. In total, these data indicate that DA levels modulate healthspan in A53T expressing animals, whereby too much DA (CAT-2 overexpression) or too little (Δ*cat-2*), reduces animal survival to N2 levels (Figure 3D-F) following exposure to paraquat.

No significant changes in oxidative stress susceptibility in either A53T(125-9m) animals or in A53T(125-9m) α-syn animals with the Δ*cat-2* mutation were observed (Figure 3D-F). On day 14, A53T(125-9m) α-syn + CAT-2 overexpression animals displayed a significant improvement in survival response following exposure to paraquat compared to N2 control (p = 0.0322) (Figure 3F).

### Modulating DA alters longevity in the α-syn A53T background

To correlate healthspan effects with aging, we next examined lifespan of all strains. Our initial analysis of longevity compared the lifespan of the strains to each other. In an examination of DA biosynthesis on lifespan, CAT-2-overexpressing animals exhibited significantly reduced lifespan compared to those expressing GFP only (p = 0.0207) (Figure 4A). Interestingly, Δ*cat-2* mutant animals also displayed a reduced lifespan compared to GFP-only worms (p = 0.0240) (Figure 4A). No significant difference between CAT-2-overexpressing and Δ*cat-2*-mutant animals in the GFP background was observed (Figure 4A).

**Figure 4.**
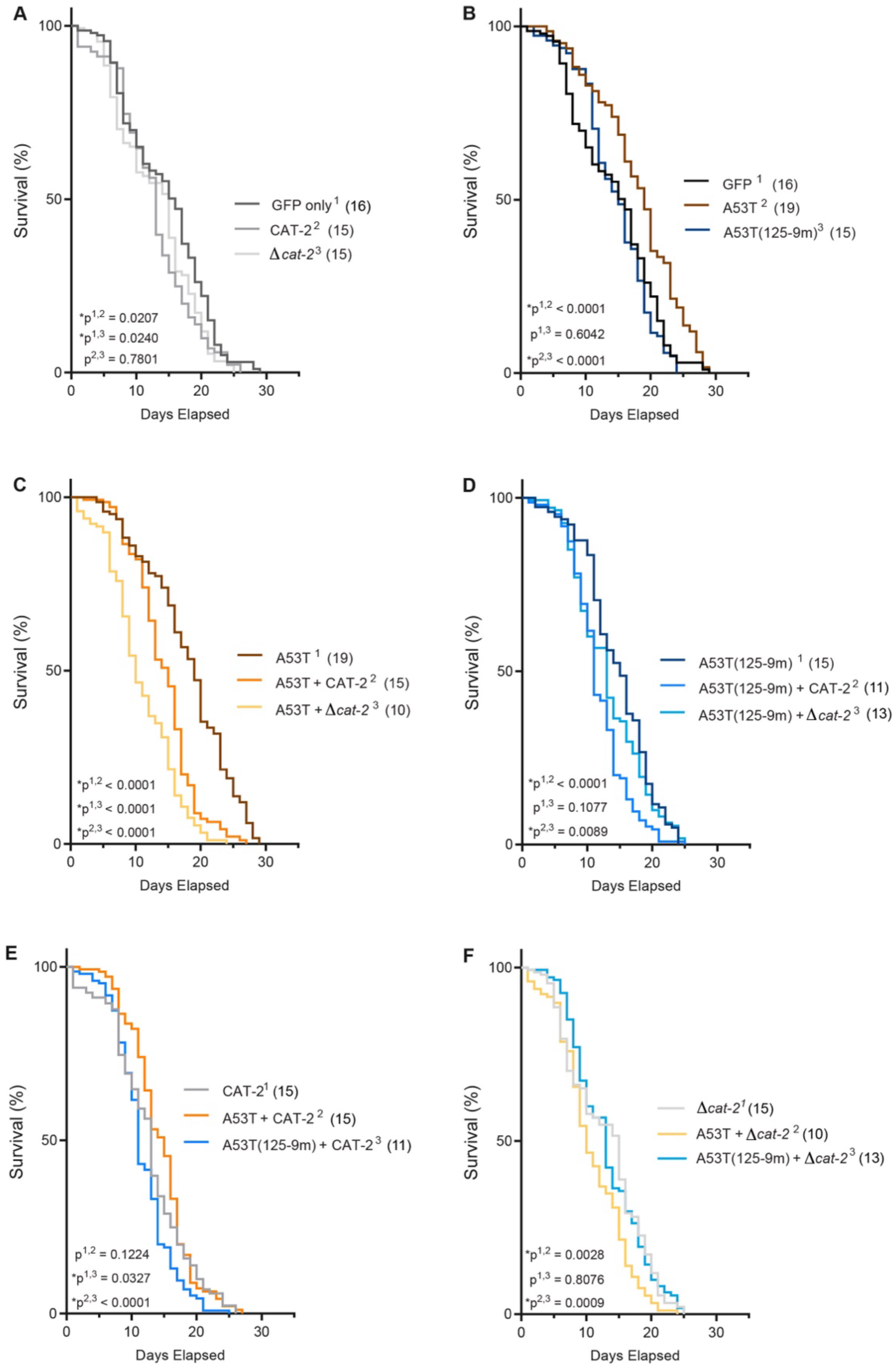
**A53T lifespan extension is dependent on basal DA levels**. **A**. Lifespan comparisons among GFP-only animals and with CAT-2 overexpression and Δ*cat-2* mutant backgrounds, where the lifespan extension was lost. Median lifespans are noted in parentheses. **B**. Animals overexpressing A53T show an extended lifespan compared to GFP only animals, where A53T(125-9M) did not. **C**. CAT-2 overexpression decreased lifespan in A53T animals. The Δ*cat-2* mutant background significantly lessened A53T lifespan compared to CAT-2 overexpression. **D**. In the A53T(125-9M) background, addition of CAT-2 overexpression resulted in a decrease of lifespan. Δ*cat-2* did not significant change in lifespan. **E**. Lifespan comparisons within the CAT-2 overexpression background demonstrate that A53T(125-9m) + CAT-2 animals live significantly shorter than A53T + CAT-2 animals. **F**. Lifespan comparisons within the Δ*cat-2* background demonstrate that A53T + Δ*cat-2* animals possess the shortest lifespan. All animals express P*_dat-1_*::GFP. n=15 animals per genotype per replicate with 10 independent replicates; total of 150 animals per genotype. Statistical differences were obtained using the Log Rank (Mantel-Cox) test.

A significant extension of lifespan was observed in A53T-overexpressing animals compared to worms completely lacking α-syn (GFP only) (p < 0.0001) (Figure 4B).

Increased DA biosynthesis from CAT-2 overexpression in the A53T background significantly reduced the lifespan extension observed in the A53T only background (p < 0.0001) (Figure 4C). Moreover, A53T animals with the Δ*cat-2* mutation revealed an even greater reduction in lifespan compared to A53T (p < 0.0001) (Figure 4C). Thus, either enhancement or depletion of DA biosynthesis negatively impacted lifespan in the A53T background. We examined these lifespan data within the context of DA levels (Figures 4E, F). There was no difference between CAT-2 expressing animals and A53T + CAT-2 (p = 0.1224). However, there was a genetic interaction between A53T + Δ*cat-2* and Δ*cat-2*, where the combination of A53T + Δ*cat-2* resulted in the shortest lifespans (p = 0.0028).

Lifespan in A53T(125-9m) expressing animals was similar to GFP only (p = 0.6042) and significantly reduced vs. A53T expressing animals (p < 0.0001) (Figure 4B). The lack of DA production in the A53T(125-9m) + Δ*cat-2* background did not impact overall lifespan when compared to A53T(125-9m) alone (p = 0.1077) (Figure 4D). While the A53T(125-9m) + Δ*cat-2* animals did not exhibit a significantly different lifespan compared to Δ*cat-2* mutants alone (p = 0.8076), they were significantly longer lived than the very short lived A53T + Δ*cat-2* animals (0.0009) (Figure 4F). CAT-2 overexpression in A53T(125-9m) animals was also examined. This background significantly reduced lifespan compared to A53T(125-9m) (p < 0.0001) (Figure 4D). These A53T(125-9m) + CAT-2 animals also displayed a significantly reduced lifespan when compared to CAT-2 alone (p = 0.0327) and A53T + CAT (p < 0.0001) (Figure 4E). Taken together, these data suggest that the changes in lifespan reflect the interactions between distinct A53T variants and DA abundance. To continue exploring the effects of aging on these complex DA-α-syn genotype interactions, we modulated autophagy genetically (Figure 5) and chemically (Figure 6).

**Figure 5.**
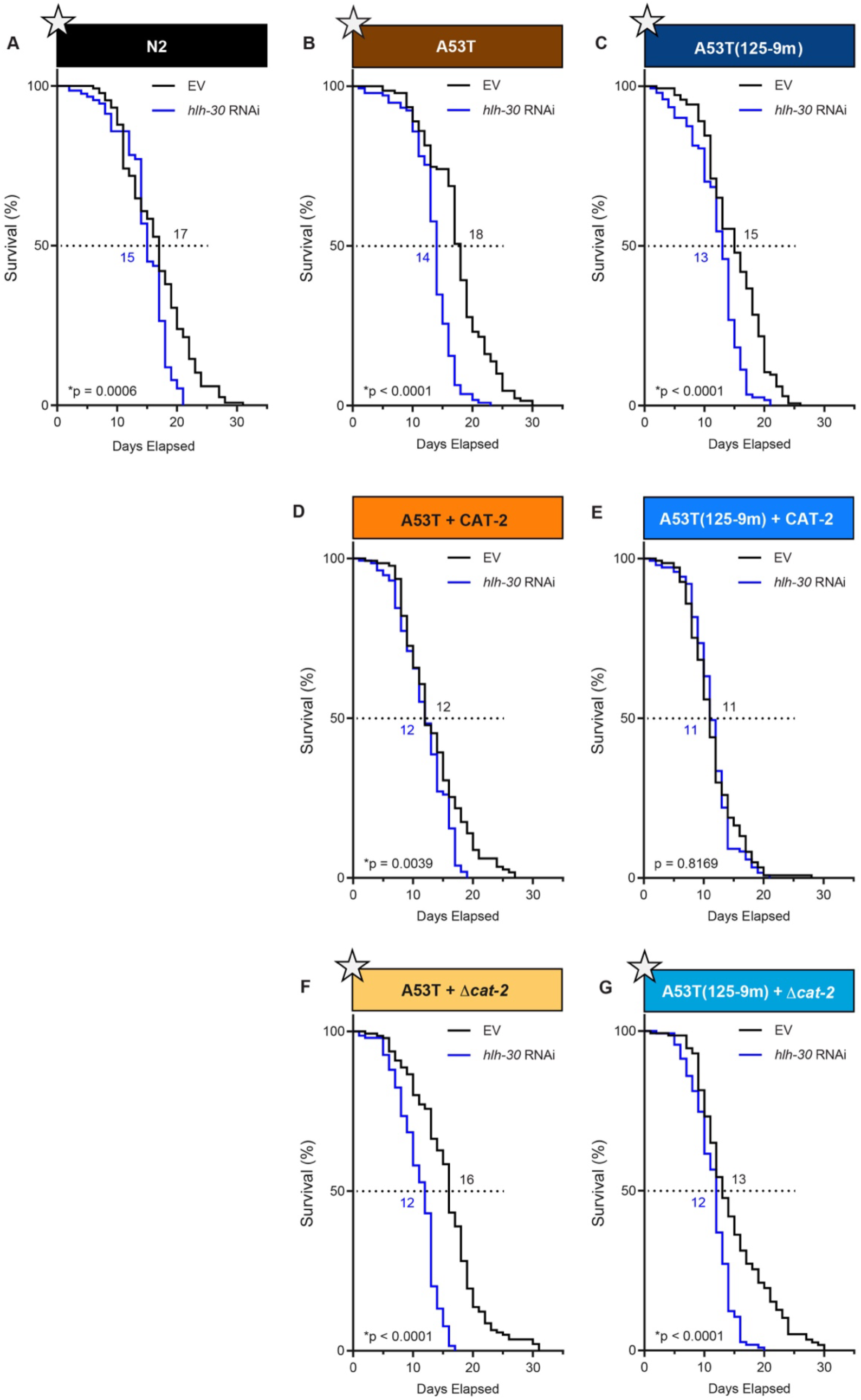
**HLH-30 is required for lifespan extension in A53T background except when CAT-2 is overexpressed**. **A-D, F, G**. RNAi of *hlh-30* in most genetic backgrounds, apart from A53T(125-9M) + CAT-2 (**E**) resulted in a significant reduction of overall lifespan when compared to EV controls. **D, E.** Both strains overexpressing CAT are non-significantly differently from their EV RNAi controls for median lifespan, while all other strains exhibited reduced median lifespan. Shortened median lifespans are emphasized with star symbols. The most dramatic decreases in median lifespan were in A53T (18 to 14 days) and A53T + Δ*cat-2* animals (16 to 12 days). n=15 animals per genotype per replicate with 10 independent replicates; total of 150 animals per genotype. Statistical differences were obtained using the Log Rank (Mantel-Cox) test.

**Figure 6.**
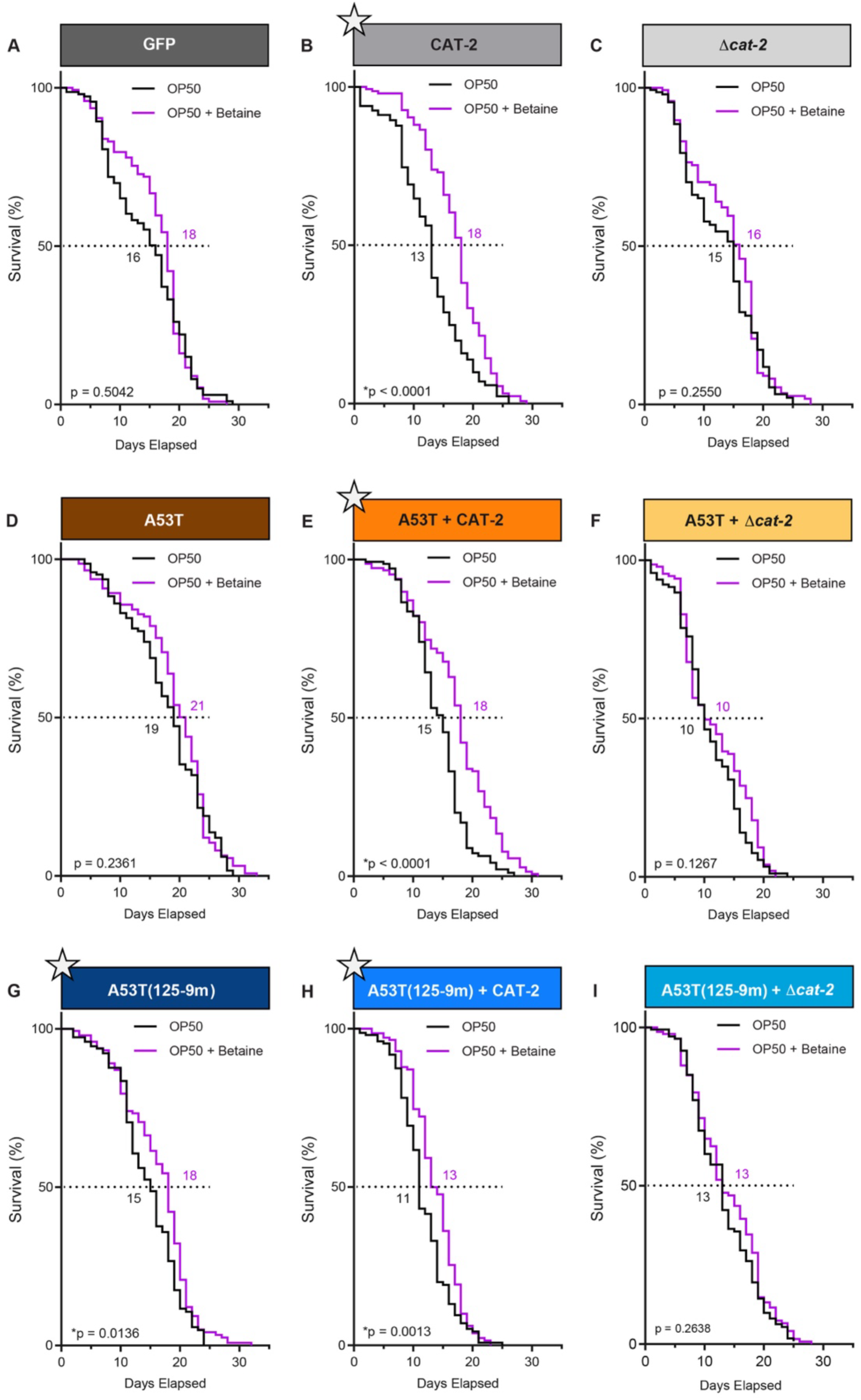
**Betaine treatment extends lifespan in a dopamine-dependent manner**. **A, C, D, F, I**. These strains are not responsive to lifespan extension from 100 µM betaine exposure. **B, E, H**. Lifelong exposure with 100 µM betaine supplementation in strains overexpressing CAT-2 resulted in lifespan extension (emphasized with star symbols). **G**. A53T(125-9m) was the only other strain that responded to betaine supplementation with lifespan extension. n=15 animals per genotype per replicate with 10 independent replicates; total of 150 animals per genotype. Statistical differences were obtained using the Log Rank (Mantel-Cox) test. Median lifespans are also noted.

### HLH-30/TFEB knockdown in A53T alters lifespan in a DA-dependent manner

In *C. elegans*, autophagy is regulated by the transcription factor EB (TFEB) homolog, HLH-30. Upon translocation to the nucleus, HLH-30 induces the transcription of autophagy-related and lysosomal genes and was shown to be required for lifespan extension across six distinct models of longevity (Lapierre et al., 2013). Given the established role of HLH-30 in autophagy-dependent longevity, we next tested whether A53T-associated lifespan extension required HLH-30 activity by knocking down *hlh-30* using RNA interference (RNAi). Because RNAi experiments require the HT115 *E. coli* strain, which independently alters *C. elegans* lifespan compared to the standard OP50 diet, direct statistical comparisons between Figure 4 and Figure 5 are inappropriate (Maier et al., 2010). However, the relative lifespan trends between genotypes within the HT115 RNAi paradigm remain highly robust

RNAi knockdown of *hlh-30* in N2 wildtype animals led to an expected shortening of lifespan. This was observed when comparing empty vector (EV) control (RNAi) to *hlh-30* (RNAi) animals (p = 0.0006). The A53T and A53T(125-9m) strains also experienced significant lifespan reduction under *hlh-30* (RNAi) conditions (Figures 5B, C) (p < 0.0001 for both). The A53T + CAT-2 strain treated with *hlh-30* (RNAi) displayed a significant lifespan reduction (p = 0.0039) (Figure 5D), however, the A53T(125-9m) + CAT-2 with *hlh-30* (RNAi) did not display a lifespan reduction (p = 0.8169) (Figure 5E). When *hlh-30* was knocked down in strains carrying the Δ*cat-2* mutation, there was a significant difference in lifespan when compared to EV for both A53T and A53T(125-9m) strains (Figures 5F, G). Taken together, these data highlight the influence of DA on HLH-30-dependent lifespan extension in A53T animals.

### Betaine enhances lifespan when CAT-2 is overexpressed

To further examine autophagy and lysosomal function as a mechanism for modulating lifespan in our A53T-DA variant strains, we utilized betaine (trimethylglycine), a senolytic known to extend *C. elegans* lifespan by promoting autophagy and antioxidant activity (Lan et al., 2024). Betaine is abundantly available in diverse animals, plants and microorganisms, where it functions as an osmoprotectant and methyl group donor. Here, we examined the impact of exogenous betaine treatment on A53T-associated lifespan extension. Betaine, at 100 µM, was incorporated into nematode growth media, for exposure throughout animal lifespan.

Betaine did not increase lifespan in GFP only animals compared to control (p = 0.5042) (Figure 6A). However, CAT-2-overexpressing animals (without A53T, but with GFP) displayed significantly enhanced lifespan compared to their solvent only control (p < 0.0001) (Figure 6B). Conversely, Δ*cat-2* mutants (without A53T, but with GFP) did not display an improved lifespan following betaine exposure compared to their solvent only controls (p = 0.2550) (Figure 6C). In α-syn A53T-overexpressing animals, exposure to betaine had no effect on lifespan extension in comparison to solvent controls only (p = .2361) (Figure 6D). Similarly, betaine exposure did not alter lifespan in A53T + Δ*cat-2* animals (p = 0.1267) (Figure 6F). Conversely, betaine exposure significantly enhanced the lifespan of A53T + CAT-2-overexpression animals compared to solvent control (p < 0.0001); (Figure 6E).

Betaine-induced lifespan extension was observed in both A53T(125-9m) expressing animals (p = 0.0136) (Figure 6G) and in A53T(125-9m) + CAT-2 animals (p = 0.0013) (Figure 6H). However, A53T(125-9m) + Δ*cat-2* animals did not display a difference in lifespan compared to solvent control (p = 0.2638) (Figure 6I). In total, these data indicate that betaine, a molecule that promotes autophagy, increased lifespan in both A53T and A53T(125-9m) α-syn animals expressing CAT-2. These data complement the *hlh-30* (RNAi) results and indicate that autophagy is correlated with DA-A53T α-syn-associated effects on longevity.

### Betaine is neuroprotective in α-syn A53T background

To determine whether betaine-mediated autophagic stimulation confers neuroprotection, we exposed animals expressing A53T in DAergic neurons to 100 µM betaine from hatching until analysis. Animals treated with solvent did not display DAergic neurodegeneration; neither did betaine-treated animals (p = 0.6640) (Figure 7A). Animals expressing A53T demonstrated significant DAergic degeneration of neuronal cell bodies and processes in the presence of solvent (Figure 7A), where only 36% of the population at day 7 displayed normal DAergic neurons. Notably, these A53T animals were significantly rescued with betaine treatment, with 61% of the population exhibiting the full, normal complement of DAergic neurons (p = 0.0043). A53T + Δ*cat-2* animals were not rescued from betaine treatment (Figure 7A; p = 0.6637). Taken together, these data suggest that betaine promotes lysosomal function and clearing of cellular waste by improving A53T proteostasis in a manner akin to TFEB/*hlh-30*.

**Figure 7.**
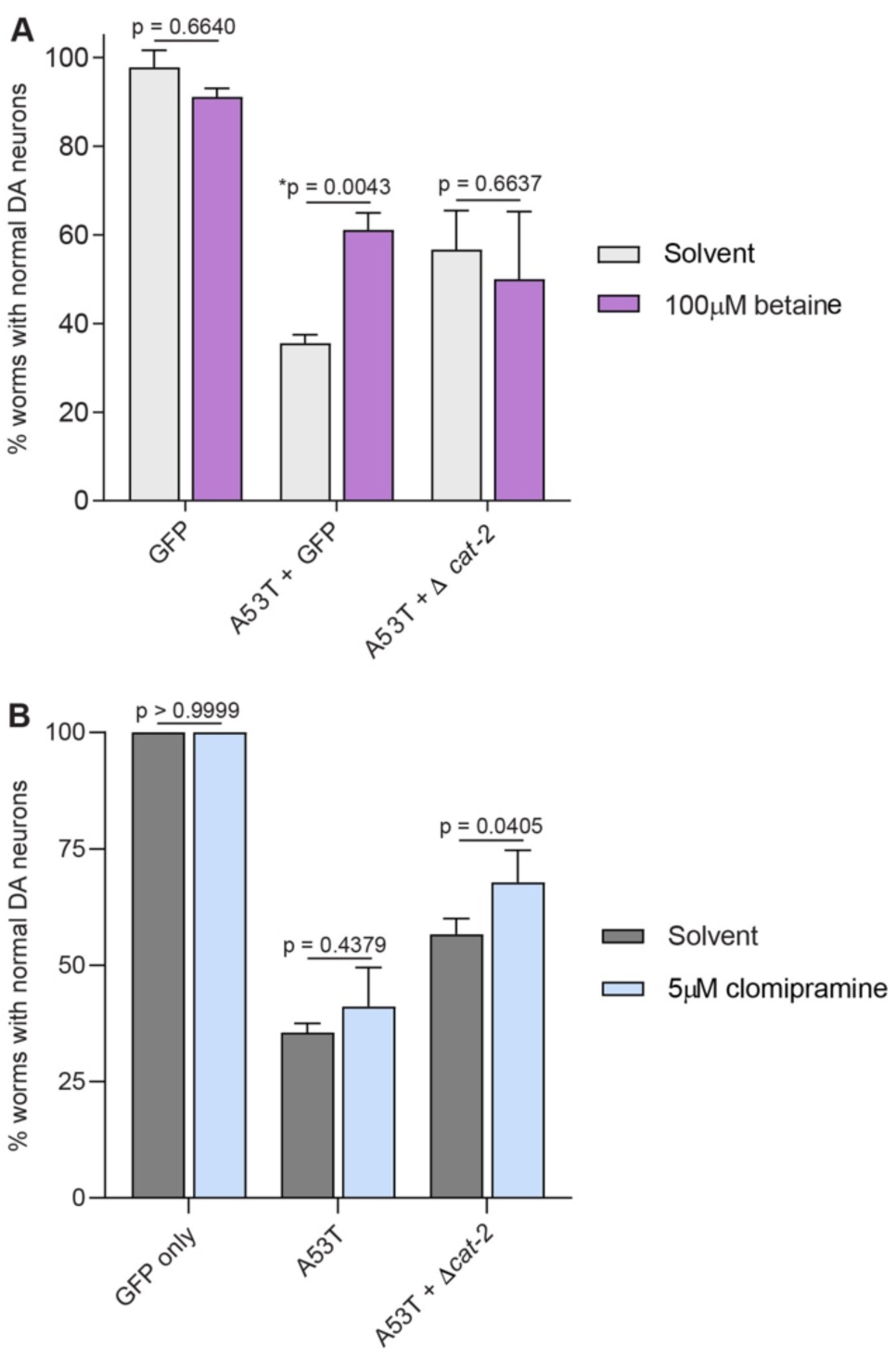
Pharmacological modulation of autophagy and lysosomal function provides context-dependent neuroprotection. **A**. Percentage of normal DAergic neurons in day 7 animals supplemented with 100 µM betaine. Betaine confers significant neuroprotection to A53T animals, but this effect is abolished in A53T animals lacking DA (Δ*cat-2*). **B**. Percentage of normal DAergic neurons in day 7 animals treated with 5 µM clomipramine. Clomipramine provides significant neuroprotection specifically to A53T + Δ*cat-2* animals. Values represent mean ± S.D. for neurodegeneration assays (n=30 worms per genotype per replicate, three replicates). Statistical significance was determined using a two-way ANOVA with Sidak’s multiple comparisons test comparing treated animals to their respective solvent-only controls.

**Figure 8.**
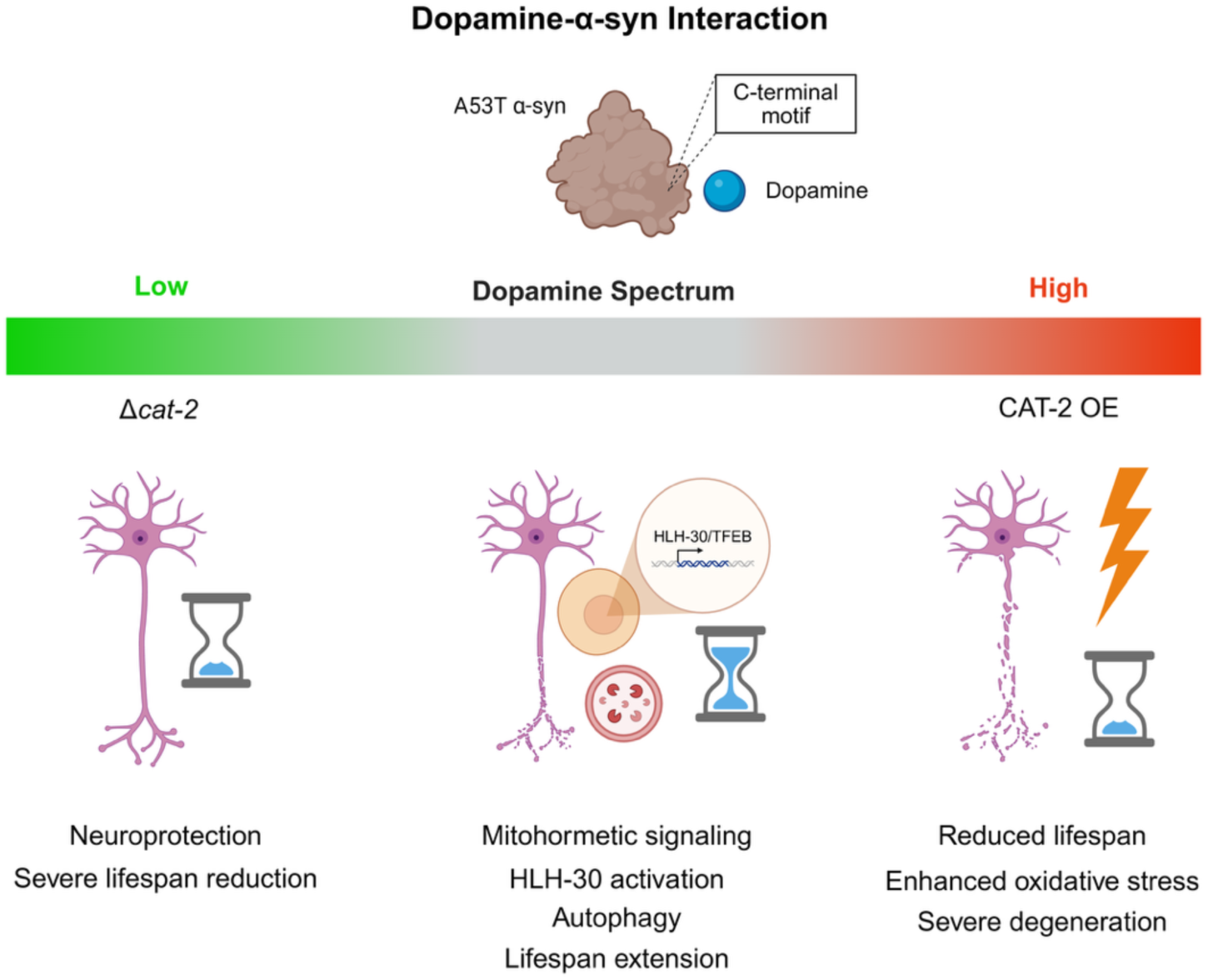
**The DA-α-syn interaction acts as a molecular mediator uncoupling neurodegeneration from systemic lifespan**. At endogenous DA levels (center), the structural interaction between DA and A53T α-syn generates a localized mitohormetic stress signal, driving moderate neurodegeneration while activating HLH-30/TFEB-dependent autophagic pathways to extend organismal lifespan. Disrupting this balance collapses the adaptive window. DA depletion (left) halts the hormetic stress response, resulting in neuroprotection but a severe reduction in lifespan, whereas excessive DA (right) overwhelms systemic proteostasis, leading to severe oxidative stress, enhanced degeneration, and a reduced lifespan. Created in BioRender. Willicott, C. (2026) https://BioRender.com/m0c44zs.

### Clomipramine is neuroprotective in the A53T + Δ*cat-2* background

A class of DA transporter (DAT) modulators that simultaneously modulate both DAT and lysosomal activity are DILAPs (DAT inhibitors and lysosomal activity promoters) (Yin et al., 2023). One of these is clomipramine hydrochloride, a drug commonly used for treatment of Obsessive-Compulsive Disorder (OCD). As with betaine, we first exposed worms expressing only GFP in DAergic neurons to 5 µM clomipramine from hatching until the day of analysis. Animals treated with solvent did not display DAergic neurodegeneration and neither did clomipramine-treated animals (p > 0.9999) (Figure 7B). Next, we examined animals expressing A53T and found that the amount of DAergic neurodegeneration was unchanged between solvent and clomipramine treatments (p = 0.4379) (Figure 7B). Notably, DAergic neurodegeneration in A53T + Δ*cat-2* animals was rescued by clomipramine treatment (p = 0.0405). Animals with normal DAergic neurons increased significantly to 68% from 57% (Figure 7B). These data suggest that the promotion of lysosomal function by clomipramine in A53T + Δ*cat-2* animals is neuroprotective, irrespective of DA depletion.

## Discussion

This study uncovered an unexpected dissociation between DAergic neuronal vulnerability and organismal aging. Prior work demonstrated that neurodegeneration in *C. elegans* synucleinopathy models is age-dependent but aging-independent, that is, the progressive degeneration of DA neurons is uncoupled from lifespan (Apfeld and Fontana, 2017).Our data identify structural DA-α-syn coupling as a pleiotropic trigger governing these distinct outcomes. Overexpression of the rate-limiting enzyme in DA biosynthesis, CAT-2 (tyrosine hydroxylase), increased DAergic degeneration in animals expressing GFP only and in the A53T α-syn background, consistent with enhanced oxidative burden (Figure 1A, G). Conversely, loss of DA synthesis through *cat-2* mutation significantly restored neuronal survival in A53T animals (Figure 1G). Mutation of the DA-interaction motif within A53T(125-9m) rendered neurodegeneration largely immutable to DA manipulation, supporting a direct structural contribution of DA-α-syn coupling to neurotoxicity in vivo (Figure 1K). These data extend prior biochemical observations demonstrating DA-mediated stabilization of α-syn oligomers and support the idea that DA enhances α-syn proteotoxicity through oxidative and conformational mechanisms (Conway et al., 2001; Norris et al., 2005; Mazzulli et al., 2007).

Strikingly, with endogenous levels of DA, A53T α-syn expression significantly extended lifespan, despite driving robust DAergic neurodegeneration. When DA was depleted in Δ*cat-2* animals, A53T α-syn expression resulted in significantly shorter-lived animals with a median lifespan of 10 days (Figure 4C); these animals also exhibited DAergic neuroprotection (Figure 1G). This tradeoff was not observed in strains lacking α-syn, where the median lifespan was maintained at wildtype levels of 15 days (Figure 4A), indicating that DA-dependent lifespan remodeling requires α-syn expression. These findings suggest that DA-α-syn coupling activates organismal management of systemic stress responses that promote longevity, even as localized neuronal damage progresses.

The uncoupling of neurodegeneration and lifespan aligns with the broader evolutionary framework of antagonistic pleiotropy, wherein DA-dependent food sensing and locomotor modulation enhance early-life fitness despite incurring an inevitable oxidative cost in late-life (Williams, 1957; Austad and Hoffman, 2018; Sawin et al., 2000). Our results further extend this paradigm by demonstrating that this localized oxidative cost actively drives systemic survival (Dues et al., 2016). In this regard, the DA-α-syn interaction functions as a mitohormetic stressor where, at endogenous levels, it generates a localized oxidative burden sufficient to trigger *hlh-30*-dependent autophagic remodeling, conferring a systemic survival advantage (Ristow and Zarse, 2010). Specifically, A53T animals exhibit an energetic advantage (Figure 3A-C) and significantly increased survival times following exposure to mitochondrial ROS-inducing agents like paraquat (Figure 3D-F). Yet, this adaptive window is highly sensitive to the magnitude of the stress. Depleting DA (Δ*cat-2*) fails to initiate this protective hormetic response, while excessive DA (CAT-2 overexpression) collapses the adaptive window when challenged with mitochondrial stressors (Figure 3). Excess DA also reduced survival back to baseline levels (Figure 4B) and precipitated severe neurodegeneration (Figure 1G).

Autophagy appears central to this adaptive response. HLH-30, the *C. elegans* ortholog of TFEB, coordinates lysosomal biogenesis and autophagic gene expression and is required for lifespan extension in multiple paradigms, including mitochondrial dysfunction and nutrient sensing (Lapierre et al., 2013). RNAi-mediated knockdown of *hlh-30* significantly shortened lifespan in N2 animals (Figure 5A), as previously reported (Lapierre et al., 2013). Likewise, *hlh-30* knockdown also significantly shortened lifespan in A53T animals (Figure 5B), demonstrating that A53T-associated longevity requires autophagic remodeling. Furthermore, chemical modulation of these pathways yielded phenotype-specific benefits. Betaine, a methyl donor that promotes autophagy, selectively enhanced survival in animals with elevated DA synthesis (Figure 6B, E, H). In contrast, DA-deficient animals failed to benefit from betaine exposure (Figure 6C, F, I), indicating that autophagy-mediated lifespan extension depends on DA-α-syn coupling. Conversely, clomipramine, a compound reported to enhance lysosomal activity, rescued neurodegeneration specifically in A53T + Δ*cat-2* animals (Figure 7B). These results suggest that boosting lysosomal function can compensate for impaired DA signaling, and that imbalanced proteostasis determines the net outcome of DA-α-syn biochemical coupling.

Collectively, these findings support a framework in which DA-α-syn biochemical coupling influences the balance between degenerative vulnerability and mechanisms of adaptive longevity. Elevated DA enhances neuronal toxicity through increased oxidative and oligomeric α-syn-induced stress, whereas intermediate levels of DA permit the activation of HLH-30-dependent autophagy to maintain systemic proteostasis. Modulation of DA biosynthesis, via *cat-2* overexpression or genetic depletion, abolishes the longevity phenotype. Thus, DA is not merely a driver of α-syn toxicity but a determinant of whether proteostatic stress engages adaptive remodeling pathways.

Therapeutic strategies that augment lysosomal capacity without exacerbating DA-driven oxidative stress may therefore preserve adaptive benefits while limiting neuronal vulnerability.

## Study limitations

One caveat of the A53T(125-9m) α-syn construct is that the substitution used to disrupt the DA-binding motif also alters residue E126 within the acidic C-terminal region of α-syn. This region has been proposed to participate in calcium binding and calcium-dependent modulation of α-syn aggregation and membrane interactions (Lautenschläger et al., 2018). Although calcium-dependent mechanisms were not examined in the present study, it remains possible that disruption of this site may influence protein conformation or aggregation dynamics independently of DA. Nevertheless, the strong correspondence between DA manipulations and phenotypic outcomes observed here supports the interpretation that DA-α-syn interactions are a primary driver of the effects described.

Additionally, while we demonstrate that A53T-associated longevity requires *hlh-30* and is responsive to lysosomal modulators, our findings rely on organismal and cellular survival phenotypes rather than direct, real-time measurements of autophagic flux or HLH-30 nuclear translocation (Zhong and Richardson, 2025). Finally, while the *C. elegans* model provides an invaluable platform to experimentally uncouple targeted neurodegeneration from organismal lifespan, it lacks the complex neuroinflammatory networks, specifically microglial and astrocytic cascades that exacerbate DAergic toxicity in the mammalian brain. Consequently, our findings isolate the cell-autonomous and systemic endocrine responses to DA-α-syn coupling. Future mammalian studies will be required to determine how this adaptive proteostatic response operates within an active neuroinflammatory environment.

## Materials and Methods

### C. elegans strains

Nematodes were maintained on NGM media with OP50 *E. coli* bacteria according to well-established laboratory conditions (Brenner, 1974). The following strains were provided by the CGC: N2, MT15620 [*cat-2(n4547)*], Integrated transgenic strain BY250 (*vtIs7* [P*_dat1_*::GFP]) was a gift from Randy Blakely (Florida Atlantic Univ.). Other strains used were MT15620 [*cat-2(n4547)*], and UA57 [*baIs4* (P*_dat-1_*::*cat-2*, P*_dat-1_*::GFP)]. Two α-syn neurodegeneration models were used in this study: UA372 (*baIs54*[P*_dat-1_*::A53T α-syn), P*_unc-54_*::tdTomato]), and UA373 [*baIs55* (P*_dat-1_*::A53T(125-9m) α-syn, P*_unc-54_*::tdTomato)]. Mutations and transgenic strains that were crossed into both UA372 and UA373 include: BY250 (*vtIs7* [P*_dat1_*::GFP]), MT15620 [*cat-2(n4547)*] and UA57 [*baIs4* (P*_dat-1_*::*cat-2*, P*_dat-1_*::GFP)]. A complete list of strains used in this study is available in Table 1.

**Table 1.** ***C. elegans* strains used in this study.**

| Strain name | Crossed strains | Genotype |
| --- | --- | --- |
| N2 | - | Bristol, wild type |
| BY250 | - | *P<sub>dat-1</sub>::GFP [vtIs7] V* |
| MT15620 | - | *cat-2(n4547)* |
| UA57 | - | *P<sub>dat-1</sub>::cat-2, P<sub>dat-1</sub>::GFP[bals4]* |
| UA372 | - | *P<sub>dat-1</sub>::α-syn(A53T), P<sub>unc-54</sub>::tdTomato[bals54]* |
| UA373 | - | *P<sub>dat-1</sub>::α-syn[A53T(125-129m)], P<sub>unc-54</sub>::tdTomato[bals55]* |
| UA502 | MT15620 x BY250 | *P<sub>dat-1</sub>::GFP [vtIs7] V; cat-2(n4547)* |
| UA525 | UA372 x UA57 | *P<sub>dat-1</sub>::α-syn(A53T), P<sub>unc-54</sub>::tdTomato[bals54]; P<sub>dat-1</sub>:: cat-2, P<sub>dat-1</sub>::GFP [bals4]* |
| UA526 | UA372 x BY250 | *P<sub>dat-1</sub>::α-syn(A53T), P<sub>unc-54</sub>::tdTomato[bals54]; P<sub>dat-1</sub>::GFP [vtIs7] V* |
| UA527 | UA372 x BY250 x MT15620 | *P<sub>dat-1</sub>::α-syn(A53T), P<sub>unc-54</sub>::tdTomato[bals54]; P<sub>dat-1</sub>::GFP [vtIs7] V; cat-2(n4547)* |
| UA528 | UA372 x MT15620 | *P<sub>dat-1</sub>::α-syn(A53T), P<sub>unc-54</sub>::tdTomato[bals54]; cat-2(n4547)* |
| UA529 | UA373 x UA57 | *P<sub>dat-1</sub>::α-syn[A53T(125-129m)], P<sub>unc-54</sub>::tdTomato[bals55]; P<sub>dat-1</sub>::cat-2, P<sub>dat-1</sub>::GFP[bals4]* |
| UA530 | UA373 x BY250 | *P<sub>dat-1</sub>::α-syn[A53T(125-129m)], P<sub>unc-54</sub>::tdTomato[bals55]; P<sub>dat-1</sub>::GFP [vtIs7] V* |
| UA531 | UA373 x BY250 x MT15620 | *P<sub>dat-1</sub>::α-syn[A53T(125-129m)], P<sub>unc-54</sub>::tdTomato[bals55]; P<sub>dat-1</sub>::GFP [vtIs7] V; cat-2(n4547)* |
| UA532 | UA373 x MT15620 | *P<sub>dat-1</sub>::α-syn[A53T(125-129m)], P<sub>unc-54</sub>::tdTomato[bals55]; cat-2(n4547)* |

### Transgenic line construction

Integrated strains UA372 [*baIs54* (P*_dat-1_*::A53T α-syn, P*_unc-54_*::tdTomato)] and UA373 [*baIs55* (P*_dat-1_*::A53T(125-9m); α-syn, P*_unc-54_*::tdTomato)] were generated from stable transgenic lines UA287 [*baEx168* (P*_dat-1_*::A53T α-syn, P*_unc-54_*::tdTomato #1); [*vtIs7* (P*_dat-1_*::GFP)] and UA288 [*baEx169* (P*_dat-1_*::A53T(125-9m) α-syn, P*_unc-54_*::tdTomato #3); *vtIs7* (P*_dat-1_*::GFP)], respectively, using a Spectrolinker XL-1500 (Spectronics Corporation, Westbury, NY, USA). Prior to integration, UA287 and UA288 were backcrossed with wildtype (N2) worms to remove P*_dat-1_*::GFP from the background. The *cat-2(n4547)* 1010 bp deletion mutation was identified during crosses using the following primers: forward GGAATAGGAACCATAGAAGATCTC and reverse CGAGGTTCTTCCTCCTAGAAATTG. The wildtype PCR product is 1208 bp and the mutant is 198 bp.

### DA neurodegeneration analysis

Animals were scored for neurodegeneration as previously described (Berkowitz, 2008). Briefly, worms were age-synchronized at hatching on NGM plates seeded with OP50 *E. coli*. All worms were kept at 20°C for the duration of each experiment. On each day of analysis, 30-35 worms were transferred to a 6 µL drop of 10 mM levamisole on a glass cover slip. The cover slip was inverted and mounted onto a 2% agarose pad on a microscope slide. Three total replicates of 30 animals were scored for each strain (3 replicates x 30 animals = 90). Using a Nikon Eclipse E600 epifluorescence microscope with the addition of a Nikon Intensilight C-HGFI fluorescent light source, the 6 DA neurons in the head region of worms (4 CEPs and 2 ADEs) were scored for the extent of neurodegeneration by using GFP fluorescence as a proxy. Each worm was scored as normal if all 6 DA neurons were present. If ADE and/or CEP neurons displayed missing processes, blebbing, and/or missing cell bodies indicative of degenerative phenotypes, then these were noted. Worms were scored for neurodegeneration on day 7 post hatching. GraphPad Prism software (version 10.6.1) was used to determine statistical significance between groups as described within individual panels of figure legends.

### Basal slowing response assays

Basal slowing response (BSR) assays were completed in a similar manner as reported by Gaeta et al (2022). Briefly, animals were grown on OP50 *E. coli* lawns and were analyzed on day 4 post-hatching. N2 and MT15620 [*cat-2(n4547)*] were used as controls. N2 exhibits a normal BSR, and Δ*cat-2* mutants display a defective one. NGM plates were used to create ring plates with a 4 cm outer ring of HB101 *E. coli* that was grown to a OD_600_ of 0.6-0.7 with an unseeded 1 cm inner circle. Each worm was allowed to briefly thrash in a 10 µL drop of 0.5x S. basal buffer to remove native bacteria. The worm was then transferred to the center of the unseeded circle of a plate at which time the video recording started. Crawling was recorded using an automated video tracking system (MBF Bioscience) and analyzed using WormLab Software (Version 4.0.5; MBF Bioscience). Each worm was recorded individually on separate plates. The average peristaltic speed (µm/sec) was recorded for the first 100 frames before and after the head of the animal touched the bacterial boundary on the unseeded portion of the ring plate. Three replicates were performed with each strain with each replicate consisting of ten individual worms. The average peristaltic speeds on and off food were compared and converted to a ratio (on/off) that was subsequently inverted and normalized to the N2 value. The resulting values represented BSR as a percentage response compared to N2 (defined as 100%). Statistical significance between groups was determined by comparing to the untreated control for each strain using an unpaired Student’s *t*-test within GraphPad Prism Software (version 10.6.1).

### Exogenous DA administration

DA plates were made by first dissolving DA hydrochloride (Fisher Scientific Cat. No. AAA1113606) in molecular grade 200 proof ethanol to a stock concentration of 500 mM and was filter sterilized using a 0.2 µm filter. Using a working stock concentration of 250 mM, 32 µL were added to each plate resulting in a final concentration of 2 mM. The plates were allowed to dry in a dark chemical fume hood for one hour and were used immediately to avoid oxidation. Control (solvent) plates contained the same concentration of ethanol as the dopamine plates but without the addition of dopamine. All plates were seeded with OP50 *E. coli* prior to the addition of dopamine. Young adult hermaphrodites (72 hours post egg lay) were transferred to dopamine and control plates and were incubated at 20°C for one hour prior to analysis. Replicates were staggered to avoid overexposure. Statistical significance between treated and untreated controls was determined using one-way ANOVA with Dunnett’s post hoc analysis. All statistical analyses were completed using GraphPad Prism Software (version 10.6.1).

### Lifespan analysis

Lifespan was measured using a variation of the Cohort survival assay (Sutphin and Kaeberlein, 2009; Amrit et al., 2014). Age-synchronized animals of each strain were prepared by a 2–3-hour egg lay with 25-30 gravid adults on NGM plates seeded with OP50 *E. coli*. At 72 hours post egg lay, 15 worms were transferred to 10 seeded plates for each strain (150 worms total per strain) representing day 1 of analysis. Worms were grown at 20°C and monitored at the same time daily until all worms were dead. Animals were transferred to fresh plates daily for the first 10 days and every other day after that. Each worm death was assigned a value of “1” and censored events (bagging, vulva protrusion, burrowing, etc.) were assigned a “0”. Statistical significance was determined using Log Rank (Mantel-Cox) test to compare survival data of all strains. All statistical analyses were completed using GraphPad Prism Software (version 10.6.1).

### Healthspan assays

*C. elegans* strain healthspan assays were determined using oxidative stress paradigms, including paraquat-induced ROS exposure and sodium azide-based immobilization for survival scoring, as previously described (Senchuk et al., 2017; Park et al., 2017). For paraquat-induced stress assays, age-synchronized experimental animals were prepared by 3-hour egg lays with 40-50 gravid young adults on five NGM plates seeded with OP50 *E. coli* per strain analyzed. At three days post hatching, animals were transferred to fresh plates to prevent starvation. Animals were analyzed at day 4, 9, and 14 post egg lay. On analysis day, individual worms were transferred to a 96-well plate with each well containing 100 µL of 40 mM paraquat dissolved in M9 buffer. Every hour, worms were analyzed for survival: if the worms were straightened out and did not respond to bright light or three soft touches, both anteriorly and posteriorly, using an eye-lash pick, they were counted as dead. The hour of death was recorded for each worm, and the median survival time was calculated for each strain on each day of analysis. Three replicates of 10 individual worms were performed with each strain. Statistical significance was determined by using one-way ANOVA with Dunnett’s post hoc test to compare strains with wild-type control. All statistical analyses were completed using GraphPad Prism Software (version 10.6.1).

For sodium azide-induced oxidative stress assay, animals were prepared in similar manner as above. Animals were analyzed at day 4, 9, and 14 post egg lay. Worms were transferred to OP50 *E. coli* seeded NGM plates containing 100 µM sodium azide four hours prior to analysis. Sodium azide plates were prepared by dissolving sodium azide in reagent grade water followed by filter sterilization using a 0.2 µm filter yielding a 10 mM stock solution. This was added directly to the liquid NGM media post-autoclave and prior to pouring the plates to achieve a final concentration of 100 µM. The thrashing frequency for each animal was quantified for each strain. Individual worms were first placed in a 10 µL drop of 1x M9 buffer to briefly thrash to remove any bacteria present and then were transferred to a 7.5 mL NGM plate that was flooded with 2 mL of 1x M9 buffer. The animal was allowed to recover for 15-20 seconds before the video recording began. The thrashing was recorded using an automated video tracking system (MBF Bioscience) and analyzed using WormLab Software (Version 4.0.5; MBF Bioscience). Each worm was recorded individually on separate plates. A one-minute video of each animal was analyzed for thrashing frequency which is defined as a change in direction of bending at the middle of the worm body. The average change in direction at middle (Hz) was converted to thrashing (bends per minute) by multiplying the frequency value by 60. Statistical significance was determined using one-way ANOVA with Dunnett’s post hoc analysis to compare strains with GFP-only control. All statistical analyses were completed using GraphPad Prism Software (version 10.6.1).

### Betaine treatment

Betaine plates were made by first dissolving betaine hydrochloride (Fisher Scientific Cat. No. A16122.30) in laboratory grade water (solvent) to a stock concentration of 100 mM and was subsequently filter sterilized using a 0.2 µm filter. An appropriate amount of stock solution was added to molten NGM media prior to pouring the plates to yield a final concentration of 100 µM. The plates were allowed to solidify before being stored at 4°C. Control plates were prepared in a similar manner without the addition of betaine. All plates were seeded with OP50 *E. coli* and incubated overnight at 37°C. Animals that were used for neurodegeneration and lifespan assays were prepped as described above with exposure starting at embryo stage. For each analyzed strain, 3 replicates of 30 animals of betaine supplementation were compared to the same amount replicates reared on NGM-only plates for neurodegeneration analysis while 10 replicates of 15 animals of betaine supplementation were compared to the same amount replicates reared on NGM-only plates for lifespan analysis. For both experiments, animals were transferred to fresh plates each day throughout the course of the experiment. Statistical significance for neurodegeneration data was determined using 2-way ANOVA with Sidak’s multiple comparisons test to compare untreated animals (solvent-only) to animals that received betaine treatment (100 µM betaine). For lifespan data significance, lifespan curves for untreated control (OP50-only) animals were compared to treated (OP50+betaine) animals using Log-rank (Mantel-Cox) test. Both analyses were completed using GraphPad Prism Software (version 10.6.1).

### Clomipramine treatment

Clomipramine plates were made by first dissolving clomipramine hydrochloride (Fisher Scientific Cat. No. AAJ6248506) in laboratory grade water (solvent) to a stock concentration of 100 mM and was filter sterilized. The appropriate amount of stock solution was added to molten NGM media prior to pouring the 5 µM clomipramine plates. The plates were allowed to solidify at room temperature prior to storage at 4°C. Control plates were prepared in a similar fashion without the addition of clomipramine to the media. All plates used were seeded with OP50 *E. coli* and incubated overnight at 37°C. Animals for neurodegeneration were prepared as described above on solvent-only and clomipramine-containing plates. For all analyses completed, animals were transferred to fresh plates each day. Statistical significance was determined using 2-way ANOVA with Sidak’s multiple comparisons test to compare untreated (solvent-only) to animals that received treatment (5 µM clomipramine). These analyses were completed using GraphPad Prism Software (version 10.6.1).

### RNA interference lifespan analysis

RNA interference (RNAi) feeding constructs were obtained from the *C. elegans* Ahringer library (Kamath et al., 2003). RNAi feeding clones empty vector (EV) and *hlh-30* were cultivated initially on LB solid media containing tetracycline (15 µg/mL) and ampicillin (100 µg/mL), and then individual colonies were grown overnight in liquid LB media containing ampicillin (100 µg/mL). IPTG was added to NGM plates for a final concentration of 1 mM, plated with RNAi feeding clones, and allowed to dry. Plates were induced overnight at 20 °C. Fertile adult hermaphrodites were allowed to lay eggs for 3-4 hours to obtain a synchronized population. Lifespan analyses as described above were completed for each strain on EV- or *hlh-30*-seeded RNAi plates. Animals were transferred to fresh plates daily for the duration of the experiment. RNAi plates were used within one week of induction and were stored at 4°C, being allowed to warm to room temperature before use. Statistical significance was determined using Log Rank (Mantel-Cox) test to compare EV- and *hlh-30-*RNAi lifespan survival curves. Median survival for each strain was also identified. All statistical analyses were completed using GraphPad Prism Software (version 10.6.1).

### RT-qPCR

Gene expression was quantified using SYBR Green-based real-time PCR. Relative expression levels were normalized using the geometric mean of multiple validated reference genes according to the geNorm method (Vandesompele et al., 2002). Briefly, at least 100 animals from UA372 and UA373 were cultivated at 20°C, collected, and total RNA was extracted using Tri Reagent (Molecular Research Center, Cincinnati, OH) according to the manufacturer’s guidelines. Genomic DNA contamination was removed from the samples using DNase I (Promega) treatment for 60 minutes at 37°C, followed by DNase Stop solution for 10 minutes at 65°C. 1µg of RNA was used for cDNA synthesis using iScript Reverse Transcription Supermix for RT-qPCR (Bio-Rad) following the manufacturer’s protocol. PCR efficiency was calculated from standard curves generated using pooled cDNA from all samples. RT-qPCR was performed using IQ-SYBR Green Supermix (Bio-Rad) with the Bio-Rad CFX96 Real-Time System. Amplification was not detected in no-template and no-reverse transcriptase controls. The target gene, α-synuclein, was measured in triplicate, and three independent biological replicates were tested for each sample. Expression levels were normalized to three reference genes (*cdc-42*, *ama-1*, and *pmp-3*) as previously described (Vandesompele et al., 2002). Normalized expression values were imported into GraphPad Prism software (version 10.6.1) for statistical analysis using the Student’s *t*-test. All primer sequences and temperatures used are described in Table 2.

**Table 2.**
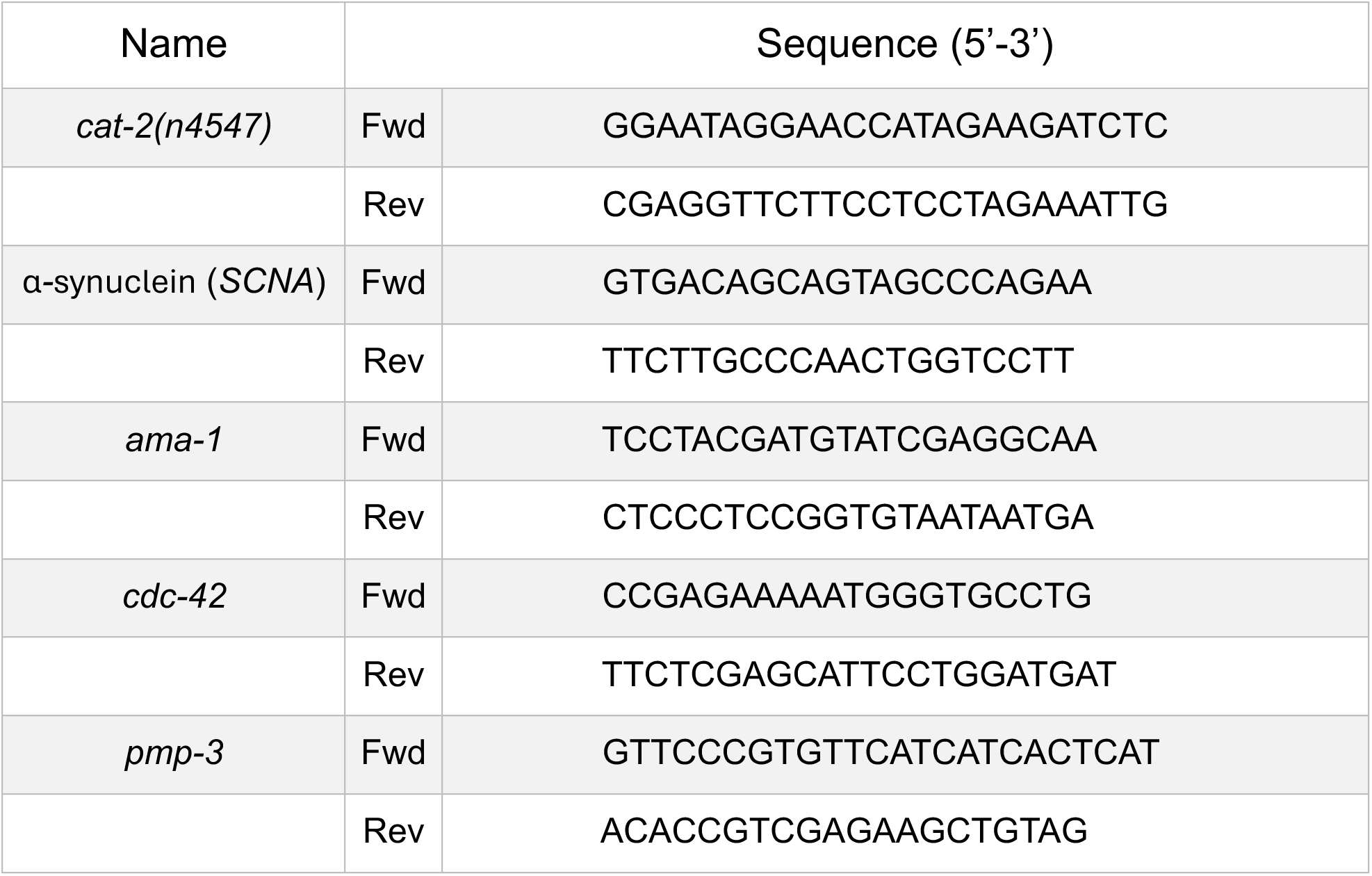
Sequences of primers for genotyping and RT-qPCR used in this study.

## Supporting information

Supplemental Info - Statistics

## Acknowledgements

Many thanks to Rachel Martin for technical assistance and the collegial environment of the entire Caldwell lab. We appreciate Randy Blakely (Florida Atlantic Univ.) for providing *C. elegans* BY250 (*vtIs7* [P*_dat1_*::GFP]). Some strains were provided by the CGC, which is funded by NIH Office of Research Infrastructure Programs (P40 OD010440).

## Data and resource availability

All relevant data and details of resources can be found with the article and its supplementary information.

## Diversity and inclusion statement

This work was a collaborative effort among multiple academic career stages, pairing a doctoral student (CWW) and undergraduate researchers (TJA, LCK) with senior mentors (LAB, GAC, KAC). This project was strengthened by the diverse, intersectional perspectives of our research team, including broad representation of geographic, generational, religious, and ability backgrounds.

